# Environmental DNA/RNA metabarcoding in estuaries of São Paulo, Brazil, reveals fish diversity and the presence of invasive species

**DOI:** 10.64898/2026.02.13.705801

**Authors:** Natascha Mozaner Nitzsche, Arthur Padial da Mota, Thomas Chen, Marcos Gomes Nogueira, Esteban Jorcin Nogueira, Naiara Guimarães Sales, Heron Oliveira Hilário, Danillo Pinhal

**Affiliations:** Laboratório Genômica e Evolução Molecular, Departamento de Genética, Microbiologia e Imunologia, Instituto de Biociências, UNESP, Botucatu, SP, Brazil; Laboratório de Ecologia de Águas Continentais, Departamento de Zoologia, Instituto de Biociências, UNESP, Botucatu, SP, Brazil; Centro de Biologia Marinha da Universidade de São Paulo, São Sebastião, SP, Brazil; School of Science, Engineering and Environment, University of Salford, Salford, UK; Laboratório de Genética da Conservação, Pontifícia Universidade Católica de Minas Gerais, Belo Horizonte, MG, Brazil

**Keywords:** Ichthyofauna, Biomonitoring, 12S rRNA, Atlantic Forest, Neotropics, Anthropogenic impact, High-throughput sequencing, Coastal conservation

## Abstract

Tropical estuaries within the Brazilian Atlantic Forest are biodiversity hotspots facing escalating anthropogenic pressures, yet their ichthyofaunal assemblages remain incompletely documented. We evaluated the combined use of environmental DNA (eDNA) and environmental RNA (eRNA) metabarcoding to characterize fish communities in two estuaries with contrasting levels of urbanization (the Juqueriquerê and Escuro rivers) on the northern coast of São Paulo, Brazil. Targeting the mitochondrial 12S rRNA (MiFish) fragment, we detected a diverse vertebrate assemblage totaling 93 species. eDNA identified 32 fish species across both systems, while eRNA detected 22 species in the preserved estuary, providing robust signals of metabolically active assemblages. The less impacted estuary exhibited significantly higher diversity indices and a more heterogeneous taxonomic composition. In contrast, the urbanized system displayed clear molecular signatures of anthropogenic influence, including the presence of invasive species (*Oreochromis niloticus, O. aureus*, and *Clarias gariepinus*) and domestic animals. This study constitutes the first application of fish eRNA metabarcoding in Brazil and demonstrates that integrating eDNA and eRNA refines ecological interpretation by coupling biodiversity detection with improved inference about contemporary community composition. Our findings highlight the potential of multi-molecule metabarcoding for routine, non-invasive biodiversity assessment in megadiverse and conservation-priority coastal ecosystems.

## INTRODUCTION

Brazilian estuaries are coastal ecosystems of extreme ecological and economic relevance, as transition areas between terrestrial and marine environments and harboring high biodiversity (Fairbridge, 1980). These environments serve as nurseries for several species of fish, crustaceans, and mollusks, playing essential roles in maintaining food chains and fish stocks (Levin et al., 2001; Zainal Abidin et al., 2022).

On the coast of the state of São Paulo in Brazil, the estuarine systems and their surroundings include several legally protected areas and are part of the largest remnants of the Atlantic Forest in Brazil (Galindo-Leal, de Gusmão Cãmara and Sayre, 2003). The Atlantic Forest Biosphere Reserve is considered a National Heritage Site by the Brazilian Federal Constitution (art. 255) and a biome threatened with extinction, including due to the degradation of natural streams and rivers, which affects many species of freshwater fish and estuarine environments (Barrella et al., 2014). It is also among the regions of the planet with the highest priority for biodiversity conservation, being considered a “hotspot” with a great wealth of species, many of which are endemic and threatened (Galindo-Leal, de Gusmão Cãmara and Sayre, 2003). The northern coast of the state is home to several estuaries and is practically entirely covered by Atlantic Forest, although human occupation has degraded 90% of its area (Malabarba, 2006; Joly et al., 2010).

Wildlife survey studies in these areas are crucial to document local diversity, identify threatened species, detect biodiversity loss and invasive species, as these ecosystems are among the most vulnerable to pollution, urban expansion, and climate change, in addition to presenting different species according to salinity variations (Barletta and Lima, 2019).

Regarding fish, the state is home to approximately 393 species of freshwater fish and 594 species of marine fish (Galindo-Leal, de Gusmão Cãmara and Sayre, 2003; Oyakawa and Menezes, 2011). However, there are gaps in species surveys in little-explored areas, such as many coastal regions, in addition to the need for frequent monitoring, especially in the Atlantic Forest areas, where the high rate of speciation and the high degree of endemism of fish communities make them even more vulnerable (Menezes, 2011; Gonçalves et al., 2017).

Different techniques are used to characterize the faunal composition of a site. Molecular identification methods in particular have been increasingly improved to provide a good margin of confidence by enabling procedures using tiny amounts of biological material, detecting target species that inhabit the ecosystem with greater precision and less effort compared to traditional field surveys (Keskin, 2014; Barreto et al., 2017; Hobbs et al., 2019).

Essentially, research has taken advantage of the high sensitivity and increasing efficiency of molecular techniques for species discrimination based on the analysis of regions of nuclear and mitochondrial DNA or the combination of both (Pank et al., 2001; Abercrombie et al., 2005; Pinhal et al., 2008, 2012; Feitosa et al., 2018), and more recently by RNA analysis (Miyata et al., 2021; Veilleux, Misutka and Glover, 2021; Yates, Derry and Cristescu, 2021).

Within the molecular approach, environmental DNA (eDNA) emerges as an indirect marker method for monitoring biodiversity, based on the genomic DNA of various organisms present in an environmental sample. EDNA may come from intracellular or extracellular sources, originating from whole microbial cells or multicellular organisms, through metabolic waste, damaged tissues or desquamated skin cells that are released into the environment (Kelly et al., 2014). From environmental samples of soil, sediment, water, feces or materials resulting from air, water or sediment filtration, it is possible to isolate eDNA with the aim of obtaining valuable taxonomic information for a given ecosystem without the need to capture an organism, an important factor when sampling vulnerable environments or species (Taberlet et al., 2018).

eDNA metabarcoding, in turn, combines universal nuclear or mitochondrial DNA barcodes (DNA barcode) and large-scale sequencing (Ambardar et al., 2016; Ruppert, Kline and Rahman, 2019; Adamo et al., 2020; Othman et al., 2023) to identify and classify organisms, allowing the simultaneous analysis of multiple sequences and producing large amounts of data on biodiversity, in less time, with good cost-benefit (Coissac, Riaz and Puillandre, 2012). The ability of eDNA metabarcoding to provide a comprehensive and rapid overview of species stands out in analyses of macrofauna and meiofauna in aquatic environments, and biomonitoring based on water analyses has gained prominence over time (Evans et al., 2016; Shaw et al., 2016; Valentini et al., 2016; Collins et al., 2019; Miya, Gotoh and Sado, 2020; Miya, 2022).

Recently, research has advanced exploring the use of environmental RNA in species identification. Although in vitro RNA has been shown to be less stable than DNA, a study with marine invertebrates demonstrated the ability to detect eRNA in water samples up to 13 hours after collection (eDNA persisted for up to 94 hours), a longer period than expected (Adamo et al., 2020). Other studies have advanced the use of eRNA and found that it can be detected from different types of RNA in the environment, as demonstrated by the analysis of miRNAs (Ikert et al., 2021), ribosomal RNA (Marshall, Vanderploeg and Chaganti, 2021; Miyata et al., 2021) and messenger RNAs (Adamo et al., 2020; Marshall, Vanderploeg and Chaganti, 2021; Tsuri et al., 2021).

Furthermore, eRNA has recently been explored as an addition to eDNA analyses, overcoming a limitation commonly faced by eDNA studies. The incidence of false positives in community analyses (target species is present, but its DNA is not recorded) and false negatives (target species is absent but its DNA is recorded) may occur when dealing with eDNA, considering that eDNA may have been transported from one location to another, preserved after the death of an animal, or moved from deeper sediments to surface layers of sediments or the water column, that is, its detection does not necessarily indicate the presence of the species (Roussel et al., 2015; Cristescu and Hebert, 2018; Miyata et al., 2021). In this context, eRNA has an advantage, considering that it degrades more quickly after cell death, reducing the chances of carryover and consequently false positives, and having the potential to be a better indicator of living biotic assemblages that are closely associated with the sampling site (Cristescu, 2019; Marshall et al., 2021).

Recent research supports eRNA as a metabarcoding analysis tool in saltwater, with results suggesting that, to unleash the full potential of ecosystem assessment, eDNA and eRNA should be studied in parallel in routine ecosystem surveys, and the degradation rates of both biomarkers assessed, thus mitigating potential false positives (Marshall, Vanderploeg and Chaganti, 2021; Miyata et al., 2021).

Evaluating eRNA is an efficient and advantageous strategy in studies aimed at analyzing the biodiversity of an estuarine community, considering that the combined use of eDNA and eRNA in environments can be advantageous for detecting rare or difficult-to-detect taxa that are not easily found by traditional methods (Stoeckle, Soboleva and Charlop-Powers, 2017; Giroux et al., 2022).

In addition, this strategy allows the identification of species in environments that are difficult to access using traditional methods and in a non-invasive manner, an important factor considering that estuarine areas function as breeding areas for many species, including those threatened with extinction. Such factors suggest that eDNA combined with eRNA can facilitate the geographic and temporal mapping of marine fish populations at a relatively low cost (Stoeckle, Soboleva and Charlop-Powers, 2017; Zainal Abidin et al., 2022). Our study aims to evaluate the combined use of eDNA and eRNA metabarcoding in the identification of fish species in two Brazilian estuaries, using eRNA metabarcoding for the first time in the country for the composition of the ichthyofauna.

## MATERIAL AND METHODS

### Study area

The eDNA and eRNA metabarcoding samples were collected in two periods of the year, August 2023 and February 2024. Our selected areas were the estuaries of two locations, the Juqueriquerê River estuary, Caraguatatuba - SP (23°42’11”S 45°26’10”W) and the Escuro River estuary region, in Praia Dura, Ubatuba - SP (23°29’24”S 45°09’57”W). The municipalities are highlighted in figure 1.

**Figure 1:**
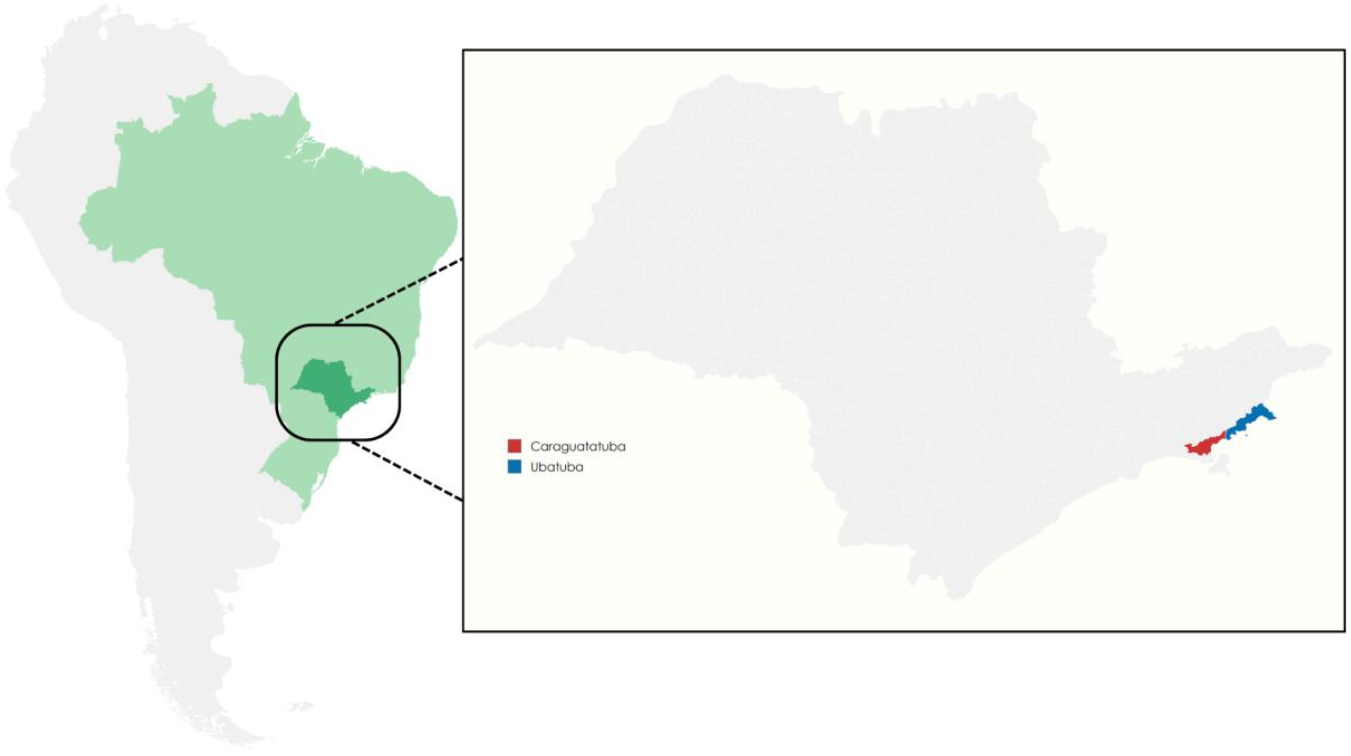
location of the municipalities chosen for collection

The Juqueriquerê River is the largest navigable river on the northern coast of São Paulo and its estuarine zone is 4.0 km long, with a semidiurnal micro-tidal regime (Rodrigues, Silva and Bernardes, 2013). The Juqueriquerê River Basin is characterized by its tourism potential and environmental problems related to human occupation and irregular land subdivision, with impacts such as the inadequate discharge of sewage and solid waste and irregular occupation of preservation areas (Lúcia et al., 2005; Boulomytis, Zuffo and Gireli, 2017; Favoretti and Batalla, 2017).

The Escuro River is located in Ubatuba - SP, which has 80% of its territory located in the Serra do Mar State Park, the largest fully protected area on the Brazilian coast. The estuarine region of this location is formed mainly by mangroves, where the Escuro and Cajarana rivers meet. In the Ubatuba region, the estuarine zone of Praia Dura has the most significant area of mangroves throughout the municipality, being part of the estuarine zones that are furthest from urbanized areas, and therefore presenting a good degree of conservation with low anthropic impact (Vasconcellos and Sanches, 2009).

### Water sampling and filtration

To collect water samples, we used 1L plastic bottles and coolers with ice to store the contents until filtration. The bottles and boxes were properly sterilized by washing them 3 times using (I) a mixture of bleach and 10% distilled water, (II) 70% alcohol and (III) 100% alcohol.

We selected 15 points along each river from the estuary mouth: (I) 5 in the mouth region, where seawater predominates, denominated as “sea”; (II) 5 in the intermediate region, where freshwater and saltwater mix, denominated as “transition”; and (III) 5 in the river region, where freshwater predominates, denominated as “river”. All samples were duplicated to cover both methodologies, totaling 30 L of water for each river. Of the total, 15 L were sent for eDNA filtration and 15 L for eRNA filtration.

All water samples for eRNA analysis were processed using SterivexTM filters (pore 0.22 μm; Millipore, USA). Disposable 50 ml luer lock syringes were attached to the filters and the volume of water from the bottle was passed through each unit using the plunger pressure. After filtration, the filters were filled with 700 μl of RNA/DNA ShieldTM (Zymo Research, Germany) and both ends sealed with parafilm (Miyata et al., 2022; Trivedi et al., 2018). All filters were stored individually in Ziplock bags or sterile falcon tubes containing silica beads, and were subsequently stored in coolers until RNA extraction.

Some samples targeted for eDNA were filtered using Sterivex filters, while the majority were filtered through 47mm diameter nitrocellulose membranes (pore 0.22μm; Millipore, USA) coupled to a vacuum pump. After filtration, the filters were folded and wrapped in currency paper, stored in zip lock plastic bags containing silica beads. The zip locks and falcon tubes containing the filters were kept in a cooler with ice until transport to the laboratory, where they remained in a freezer at -20oC until the time of genetic material extraction.

### eDNA/eRNA extraction

To extract eDNA from nitrocellulose membranes, we inserted each unit into a 1.5 ml Eppendorf tube using properly sterilized scissors and forceps. Each membrane was cut and macerated using scissors, and then 80 ul of proteinase K and 750 ul of lysis buffer were added. We performed the remaining steps of the extraction process according to the NucleoSpin® Tissue Kit protocol (Macherey Nagel, USA).

To extract eDNA and eRNA from SterivexTM filters, we arranged each filter vertically, with the larger end facing down, and supported on a 1.5 ml Eppendorf tube. Both were housed in this configuration inside a 50 ml falcon tube, ensuring the stability of the filter position. The falcon tubes were centrifuged for 1 min at 1500g, causing the RNA/DNA Shield content retained in the tube to concentrate in the Eppendorf tube. For eDNA samples, we subsequently added 180 ul of lysis buffer and 25 ul of proteinase K (NucleoSpin® Tissue Kit, Macherey Nagel, USA) and performed the remaining steps of the extraction process according to the NucleoSpin® Tissue Kit protocol (Macherey Nagel, USA). For eRNA samples, we followed the same procedure performed for eDNA to remove the RNA/DNA Shield content from the SterivexTM filters. Subsequently, we added 750 ul of lysis buffer (Quick-RNATM Miniprep Kit, Zymo Research) and followed according to the manufacturer’s protocol.

The extracted total RNA and DNA were measured directly by fluorometry (QubitTM RNA High Sensitivity (HS) and QubitTM 1X dsDNA High Sensitivity (HS), Life Technologies, USA) and the purity determined by the absorbance parameters in NanoDrop® ND-1000 (Thermo Scientific).

The eDNA samples were sent for DNAse treatment with the Turbo DNA-free kit (Thermo Fisher) following the manufacturer’s protocol. To create the cDNA library, we performed reverse transcription reactions using the High Capacity cDNA Reverse Transcription Kit (Applied Biosystems, USA).

### PCR Amplification and DNA Sequencing

The cDNA and DNA samples were amplified via PCR. Target amplicons were generated from the 12S ribosomal (rRNA) gene of mitochondrial DNA, using the universal primers MiFish-U-F/R (U, F and R represent universal, forward and reverse, respectively), already used for fish metabarcoding (Miyata et al., 2021 and Miya et al., 2015). The selected primers target approximately 170 base pairs in the region at the beginning of the 12S rRNA gene, which contains sufficient variability to allow the identification of fish to the family, genus and species level, with the exception of some closely related genera (Polanco F. et al., 2021). MiFish-E-F/R was designed to encompass sequence variations present in elasmobranchs.

The PCR was divided into two steps to prepare the amplicon library for high-throughput sequencing, following the two-step PCR-based approach described in the Illumina 16S metagenomic sequencing library (Illumina, Inc., San Diego, CA, USA), already used in 12S eDNA protocols (Evans et al., 2017; Bevilaqua et al., 2020; Kumar et al., 2022).

The reactions were initially performed using the InvitrogenTM PlatinumTM Taq DNA Polymerase kit under the following conditions: initial denaturation at 94 °C for 2 min, followed by 35 cycles of denaturation at 95 °C for 30 s, annealing at 53.8 °C for 30 s, and extension at 72 °C for 30 s, with a final extension at 72 °C for 5 min.

After completing the initial tests, we performed PCR reactions using primers with adapters and the high-fidelity DNA polymerase NEBNext® Q5® Hot Start HiFi PCR Master Mix, adapting the conditions to the manufacturer’s protocol under the following conditions: initial denaturation at 98 °C for 30 s, followed by 35 cycles of denaturation at 98 °C for 10 s and annealing at 53.8 °C for 30 s, with final extension at 65 °C for 5 min. The samples were visualized on an agarose gel and purified using AMPure XP Bead-Based Reagent metal beads, following the manufacturer’s protocol. The intact samples were sent to the second indexing stage. We used the Illumina DNA/RNA UD Indexes Set A kit for the second PCR, also performed using NEBNext® Q5® Hot Start HiFi PCR Master Mix, under the following conditions suggested in the protocol and optimized in the laboratory: initial denaturation at 98°C for 30s, followed by 14 cycles of denaturation at 98°C for 10s, annealing at 53.8°C and extension at 65°C, with a final extension at 65°C for 5 minutes. The PCR products again underwent purification using AMPure XP Bead-Based Reagent metal beads, with subsequent standardization of product concentration and integrity analysis performed using BioAnalyzer or TapeStation (Agilent Technologies, Inc., San Diego, CA, USA) (Miya et al., 2015; Miya, Gotoh and Sado, 2020).

For Illumina paired-end sequencing, we used the micro V2 300 cycles kit, following the 16S library preparation protocol provided by Illumina, and the 20 samples were inserted into the MiSeq (Illumina).

### Data analysis

The samples containing the raw sequences had undetermined bases removed and quality filtered (Q-scores ≥ 30). Only reads containing matching primers and adapters were targeted for error correction and subsequent read analysis using the DADA2 Pipeline, presented by Callahan and collaborators, with the potential to identify real variants and produce few false overestimated sequences, working with amplicon sequence variants (ASVs) to model and correct errors in amplicons sequenced by Illumina (Callahan et al., 2016; Nearing et al., 2018; Miya, Gotoh and Sado, 2020; Minamoto et al., 2021).

All identifications were made using NCBI nucleotide and NCBI taxonomy available in the NCBI database. (http://www.ncbi.nlm.nih.gov). Considering that in few samples it was possible to join the sequences by merging into unique sequences (ASVs), as the results are usually grouped, we also added concatenated analyses, associating the obtained sequences without pairing.

We used RStudio to graphically represent the main alpha diversity indices (Chao1, Shannon, Simpson), using the Phyloseq package (McMurdie and Holmes, 2015; Kandlikar et al., 2018), where index values are calculated for each sample based on abundance, with averages and outliers that allow us to compare the averages between rivers and understand the homogeneity of samples. Alpha diversity, often expressed as species richness, represents the diversity of species within an area, considering the distribution of the number of individuals per species (Sanches et al., 2012; Magurran, 2021).

## RESULTS

Due to the excessive presence of sediments, it was not possible to filter the total volume of water collected.

In 2023, for the Juqueriquerê River, the amount of water filtered for eRNA ranged from 170ml to 550ml, while for the Escuro River, where there was less sediment, the filtrate ranged from 500ml to 750ml. For the eDNA samples from the Escuro River, the filtered volume was 500ml for the samples using Sterivex™ and 1L for samples using nitrocellulose membranes. The eDNA samples from the Juqueriquerê River that were filtered with Sterivex™ ranged in volume from 170ml to 550ml.

In 2024, we found greater difficulty in filtration due to summer rains and accumulation of debris, with the amount of water filtered for eRNA in the Juqueriquerê River varying from 150ml to 350ml, while for the Escuro River, from 350ml to 850ml. The eDNA samples were all standardized for both rivers, in 500ml using nitrocellulose membranes.

We observed that the eDNA samples extracted in Ubatuba using Sterivex™ filters presented higher total eDNA concentration values, even with a lower volume of filtered water when compared to the total volume filtered under vacuum (1L).

In relation to the eDNA/eRNA comparison, the total eRNA concentrations extracted were also considerably higher in general compared to eDNA for both periods and rivers.

After PCR, 20 samples achieved successful amplification, all of them from 2023, with concentrations ranging from 58.1 to 106 ng/µl.

We achieved successful sequencing in 18 samples, of which 10 eDNA samples from the Juqueriquerê River, 5 eRNA samples from the Escuro River, and 3 eDNA samples from the Escuro River (**Table 1**).

**Table 1:** Sample identification (JQ = Juqueriquerê River; RE = Escuro River), with their respective source of genetic material, post-PCR DNA concentration, and filtered water volume.

| Sample | Source | Concentration (ng/ul) | Filtration volume (ml) |
| --- | --- | --- | --- |
| JQ01 | eDNA | 78,5 | 170 |
| JQ02 | eDNA | 74,6 | 170 |
| JQ03 | eDNA | 76,3 | 200 |
| JQ04 | eDNA | 103 | 200 |
| JQ05 | eDNA | 97,1 | 450 |
| JQ06 | eDNA | 100 | 450 |
| JQ07 | eDNA | 106 | 400 |
| JQ08 | eDNA | 83,3 | 500 |
| JQ09 | eDNA | 65,4 | 420 |
| JQ10 | eDNA | 76,7 | 450 |
| JQ12 | eDNA | 69,8 | 550 |
| RE02 | eRNA | 75,7 | 500 |
| RE03 | eRNA | 67 | 550 |
| RE04 | eRNA | 72,9 | 550 |
| RE05 | eRNA | 72,1 | 500 |
| RE06 | eRNA | 73,1 | 450 |
| RE10 | eDNA | 62,1 | 500 |
| RE15 | eDNA | 58,1 | 500 |
| RE16 | eDNA | 64,3 | 500 |

We identified 93 species in total, including fish, amphibians, mammals and birds. Considering only fish, we identified 32 species using eDNA for both rivers, while with eRNA we were able to identify 22 species, all from the Escuro River. Ten fish were identified using both methods (**Figure 2**). The fish that were identified using both eRNA and eDNA were the following species: Mojarra listrada (*Eugerres plumieri*), Guavina-do-Atlântico (*Dormitator maculatus*), Tainha (*Mugil platanus*), Robalo-branco (*Centropomus undecimalis*), Corvina (*Micropogonias furnieri*), Coridora-bandata (*Scleromystax barbatus*), Carapau (*Caranx latus*), Acará *(Geophagus brasiliensis*), Carapicu (*Eucinostomus melanopterus*) and Parati (*Mugil curema*).

**Figure 2:**
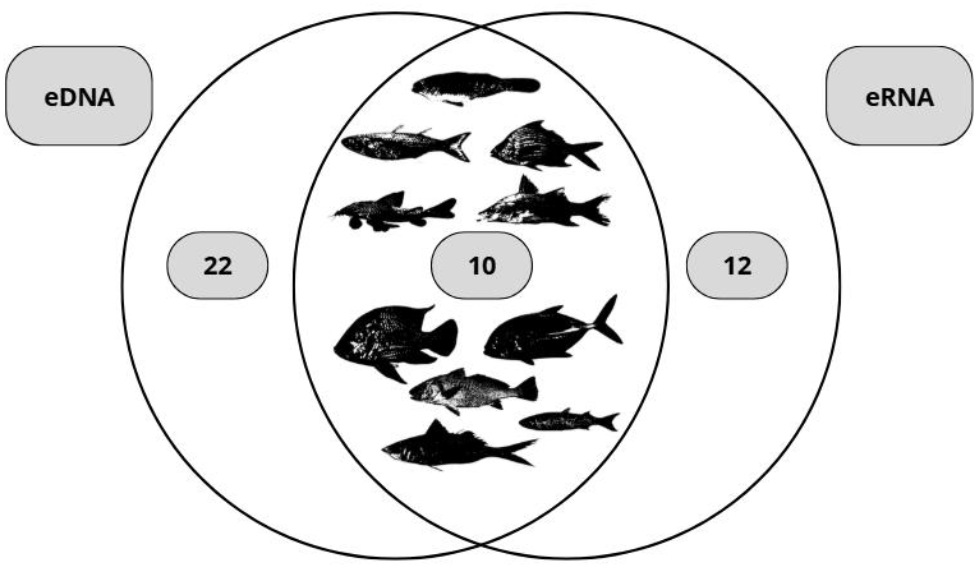
Venn diagram representing the number of species detected by each technique, respectively eDNA only (22), both (10) and eRNA only (12).

From the total number of ASVs, we infer the quantity corresponding to each percentage of identification by BLASTn, with the vast majority being identified at a percentage of 100%, allowing us to arrive at the precise identification of the species (**Figure 3**). Considering the Actinopterygii fish, we were also able to represent the richness of genera per sample and compare the diversity of information provided by each portion of the estuary in each river, demonstrating greater richness in the “river” portion of the Escuro River and in the “transition” portion of the Juqueriquerê River (**Figure 4**).

**Figure 3:**
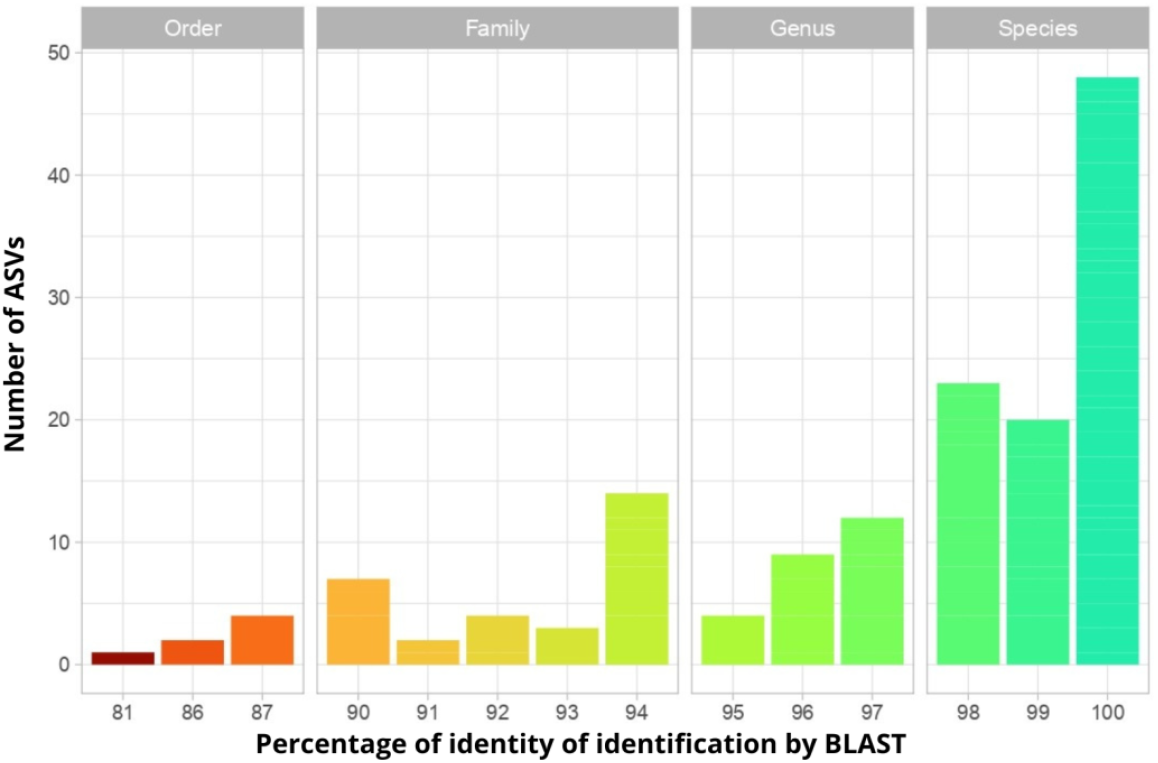
number of ASVs and their respective percentage of identification by BLAST.

**Figure 4:**
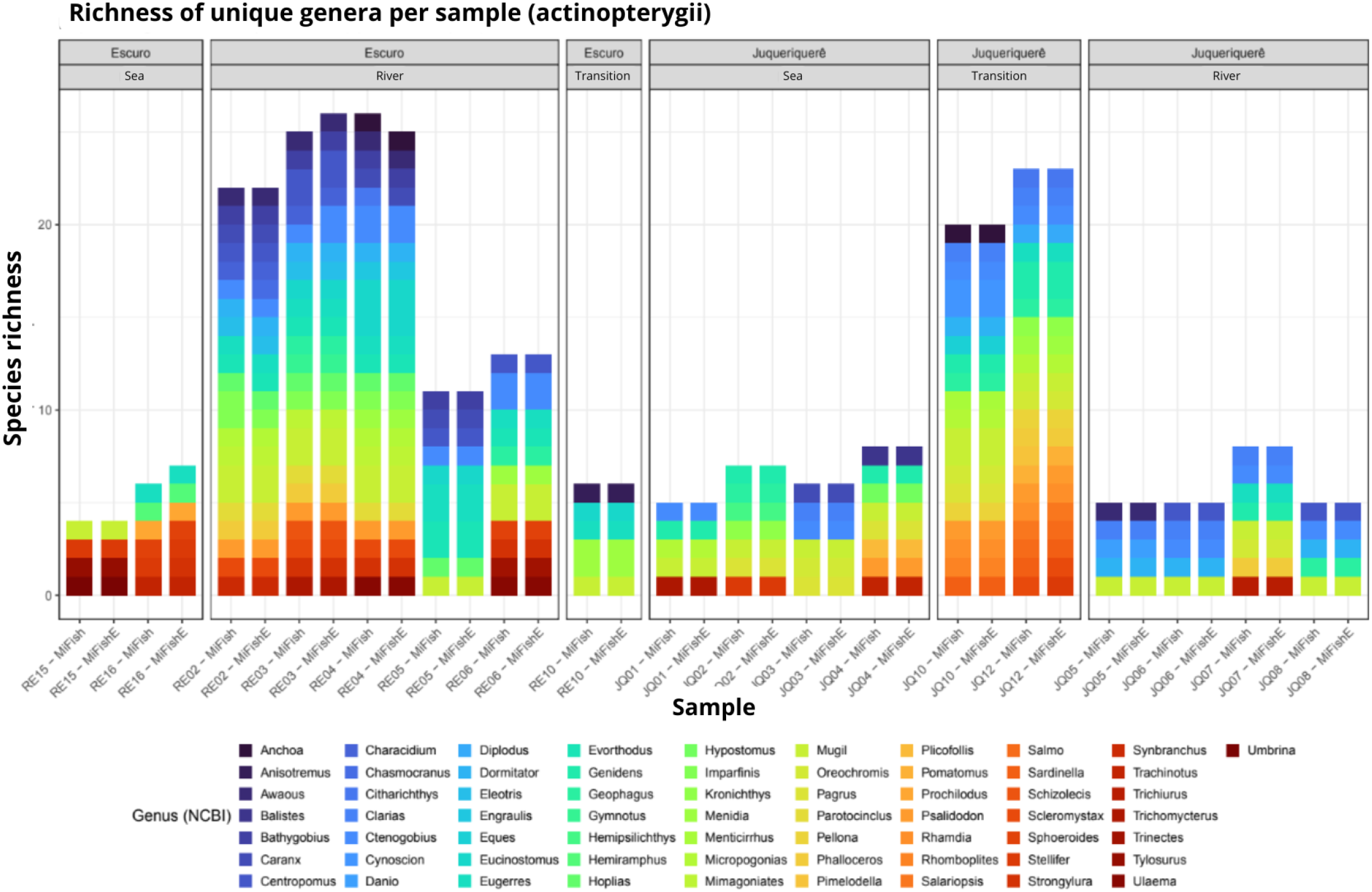
Taxon richness per sample (considering Actinopterygii)

The relative abundance of each species per sample also allows us to visualize the representation of each taxon in percentage for each of the sampled points (**Figure 5**).

**Figure 5:**
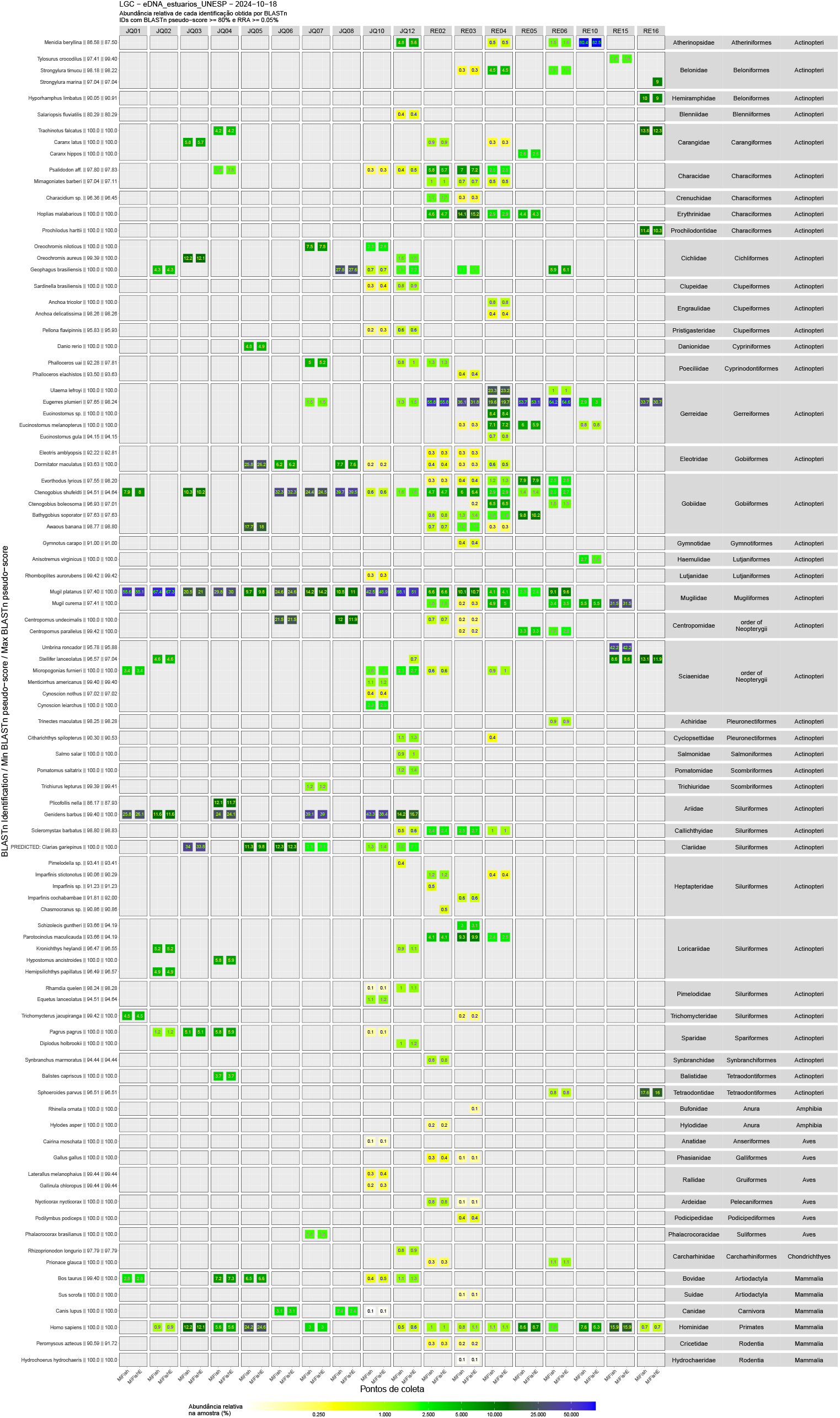
Relative abundance of each species per sample.

The main diversity indices for both rivers were compiled in boxplots (**Figure 6**). Considering the average number of species in each river, in all parameters evaluated, the Escuro River presented a greater diversity of species in relation to the Juqueriquerê River.

**Figure 6:**
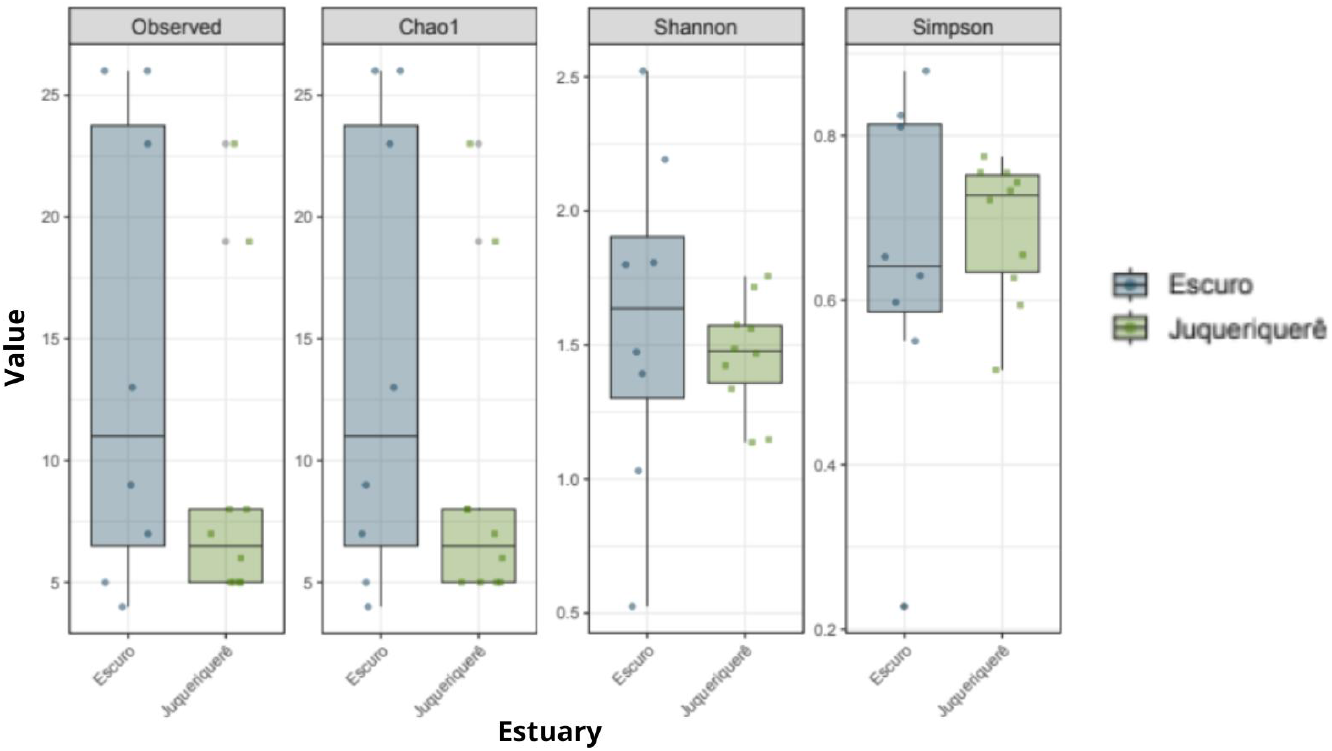
Diversity indices boxplots for both rivers, revealing higher levels of diversity for the Rio Escuro

## DISCUSSION

From the results obtained in the first stages, we noted that only the 2023 samples achieved successful amplification and ease of filtration, with no samples from 2024 being visualized in agarose gel and presenting satisfactory conditions of concentration and purity. Considering environmental factors as determinants in the availability, abundance and degradation of nucleotides in environmental samples, from the literature we assume that the intense period of summer rains in 2024 possibly contributed to the increase in water volume and dilution of genetic material, in addition to the high temperatures and turbidity caused by the movement of water, which possibly influenced the acceleration of DNA and RNA degradation (Barnes et al., 2014; Collins et al., 2018).

Considering the average number of species identified in each river, we demonstrated a high potential of eRNA in providing species identification data, considering that with only 5 successfully sequenced samples we obtained a large number of species compared to the eDNA results.

All samples from the Juqueriquerê River showed a considerable abundance of Tainha (*Mugil platanus*), which were found in smaller quantities in the Escuro River. This result is in line with expectations, since Tainha is considered one of the most abundant pelagic species in shallow coastal regions and estuaries in the state of São Paulo, and is a major target for fishing, especially artisanal fishing on the state’s coast, with most catches occurring in the winter months in Brazil (June to August), coinciding with the period in which our samples were collected (Miranda and Carneiro, 2007). The Juqueriquerê River also showed an abundance of white catfish (*Genidens barbus*), which inhabits estuarine regions and rivers, mainly in shallow and muddy bottom waters, and in Brazil occurs from the state of Bahia to Rio Grande do Sul (Ribeiro, 2011; Avigliano, Velasco and Volpedo, 2015).

We also highlight the presence of invasive fish species in the Juqueriquerê River: Nile tilapia (*Oreochromis niloticus*), blue tilapia (*Oreochromis aureus*), and African catfish (*Clarias gariepinus*). In an informal conversation between our team and local fishermen, we had already been informed of the frequent tilapia fishing in the area, which was later corroborated by our data. In recent years, there has been growing concern about the expansion of tilapia occupation in Brazil, spread mainly by aquaculture. Typically freshwater, Tilapia have recently shown potential to spread through marine ecosystems, being reported in brackish and saline waters in several regions of Brazil, including the southeast (Forneck et al., 2021; Franco et al., 2024). This situation raises concerns regarding the licensing of aquaculture projects in rivers and estuaries, since tilapia can significantly affect native species, including the acará (*Geophagus brasiliensis*), another cichlid that is quite common in river and estuary regions and was also identified in our samples, both in the Juqueriquerê and Escuro rivers. The long-term concern is that with the niche overlap between the two species and a tendency for tilapia to behave aggressively compared to the acará, there will be a reduction in the population viability of native species and competitive exclusion (Linde-Arias et al., 2008; Sanches et al., 2012). The African catfish (*Clarias gariepinus*) is also an invasive species that has spread through several Brazilian rivers and is targeted by aquaculture. In addition to being an excellent predator and competitor with native species for resources, it is a fish with a broad tolerance to anthropized environments, especially due to physiological conditions associated with the species’ respiratory tract that allow it to breathe in places with low oxygenation and poor water quality (Moreira and Silva, 2023).

When dealing with eDNA, contamination is a major challenge and can occur at all stages of laboratory protocols, including collection, extraction, amplification, and eDNA library construction, with field collections being the most critical stages of cross-contamination (Xing et al., 2022). Although all due care was taken by the team during the collection and extraction stages, including sterilization as described in protocols, the contamination risk in the laboratory environment is still high, as we found in our results (Thomsen and Willerslev, 2015; Xing et al., 2022). We observed a low abundance of zebrafish (*Danio rerio*), a research model used in our laboratory, in one of our samples, as well as blue shark (*Prionace glauca*), also with recently manipulated samples. We highlight here the importance of blank samples that signal the presence of false negatives, identifying contamination.

Non-target species were also identified, including humans and domestic animals, a result that was expected mainly in the Juqueriquerê River, with visible human occupation around it and domestic dwellings with animal husbandry. In the samples from the Juqueriquerê River, humans (*Homo sapiens*), cattle (*Bos taurus*) and domestic dogs (*Canis lupus*) were detected. The Escuro River presented, although in low abundance, the presence of domestic birds (*Gallus gallus*) and pigs (*Sus scrofa*), due to some local settlements near the river.

Although the samples successfully amplified do not provide a definitive comparison between areas, river sections, and methods, mainly due to the absence of sequenced eRNA samples from the Juqueriquerê River and the low number of eDNA samples from the Escuro River, the information gathered provides a general overview of the local ichthyofauna. The study also allowed us to demonstrate the potential of methods using eDNA and eRNA for species identification and fauna surveys, providing unprecedented information on several species that would be difficult to capture with traditional sampling techniques, especially in mangrove regions. Our eRNA samples were responsible for providing a great richness of species, confirming that eRNA is sufficiently present and persistent in the environment to be a monitoring tool, highlighting its potential to be associated with eDNA and detect metabolically active organisms (Yates et al., 2021; Stevens and Parsley, 2023). We believe this is the first successful eRNA analysis performed by researchers in the country. Nevertheless, we highlight the importance of traditional survey techniques, which provided us with information on species that were not detected by environmental techniques, being essential to confirm the real occurrence of a species and avoid the detection of false positives by eDNA, in addition to overcoming the limitation faced by the use of the 12S gene in megadiverse areas that have species not represented in databases (Zainal Abidin et al., 2022). In addition, DNA extraction from tissues from fish collections allowed us to obtain reference 12S sequences that will be used to enrich the databases. Environmental parameters associated with eDNA data can also be useful in assessing levels of anthropization and their relationship with species abundance. The Juqueriquerê River, in addition to having a lower abundance of species, also showed detections of species associated with urbanized environments, demonstrating that eDNA can be used for the early detection of invasive species. Even so, this relationship between anthropization and species richness should be viewed with caution, considering that our eRNA samples showed a greater potential for species identification and were also the largest group sequenced from the Escuro River.

Additionally, the estuarine environments evaluated, although components of the Atlantic Forest, could naturally have previously presented differentiated species richness, regardless of anthropic action. Unfortunately, there are no records of previous historical data on biodiversity that would allow us to make greater previous comparisons.

Finally, the use of combined sampling techniques proves to be an ideal portrait, optimizing the disadvantages of each technique and allowing a more accurate portrait of the local faunal composition.

## CONCLUSION

From our analysis, we conclude that eRNA and eDNA have the potential to be widely used as tools for fauna monitoring. Although they have their drawbacks, such as the need for well-established databases, contamination concerns, low eRNA stability, and the limitation of eDNA in providing information on populations and status of individuals within a population, both have proven to be efficient in metabarcoding approaches, being non-invasive and allowing a large number of species to be detected from a few samples in a short time. We also conclude that the combined use of fauna survey methods reduces the limitations of each technique and accurately represents the species composition, especially with eRNA and its detection of active organisms, representing a faithful portrait of the living fauna in a given space-time.

## ACKNOWLEDGMENTS

We sincerely thank Talita Roberto Aleixo de Almeida, Beatriz Jacinto Alves Pereira and Gabriel Mariano Silva for the support throughout the project and Leonardo Nazario de Moraes for his assistance and technical support. We thank CAPES 88887.696117/2022-00, FAPESP 2023/07311-3 and CNPq 10379 for funding research projects.

